# Ontogeny and function of the circadian clock in intestinal organoids

**DOI:** 10.1101/2020.04.22.053223

**Authors:** Andrew E. Rosselot, Miri Park, Toru Matsu-Ura, Gang Wu, Danilo E. Flores, Krithika R. Subramanian, Suengwon Lee, Nambirajan Sundaram, Taylor R. Broda, Heather A. McCauley, Jennifer A. Hawkins, Kashish Chetal, Nathan Salomonis, Noah F. Shroyer, Michael A. Helmrath, James M. Wells, John B. Hogenesch, Sean R. Moore, Christian I. Hong

**Affiliations:** Department of Pharmacology & Systems Physiology, University of Cincinnati, Cincinnati, OH 45267; Division of Human Genetics and Immunobiology, Center for Chronobiology, Department of Pediatrics, Cincinnati Children’s Hospital Medical Center, Cincinnati, OH 45229; Department of Pediatric Surgery, Cincinnati Children’s Hospital Medical Center, Cincinnati, Ohio, 45229; Center for Stem Cell and Organoid Medicine, Division of Developmental Biology, Cincinnati Children’s Hospital Medical Center, Cincinnati, OH, 45229; Division of Biomedical Informatics, Cincinnati Children’s Hospital Medical Center, Cincinnati, OH, 45229; Gastroenterology and Hepatology, Baylor College of Medicine, Houston, Texas, 77030; Division of Endocrinology, Cincinnati Children’s Hospital Medical Center, Cincinnati, OH, 45229; Center for Chronobiology, Cincinnati Children’s Hospital Medical Center, Cincinnati, OH, 45229; Division of Pediatric Gastroenterology, Hepatology, and Nutrition, Department of Pediatrics, University of Virginia School of Medicine, 1400 West Main Street, Charlottesville, VA 22908; Division of Developmental Biology, Cincinnati Children’s Hospital Medical Center, Cincinnati, OH, 45229

## Abstract

Circadian rhythms regulate diverse aspects of gastrointestinal physiology ranging from the composition of microbiota to motility. However, development of the intestinal circadian clock and detailed molecular mechanisms regulating circadian physiology of the intestine remain largely unknown. The lack of appropriate human model systems that enable organ- and/or disease-specific interrogation of clock functions is a major obstacle hindering advancements of translational applications using chronotherapy. In this report, we show that both pluripotent stem cell-derived human intestinal organoids engrafted into mice and patient-derived human intestinal enteroids (HIEs) possess robust circadian rhythms, and demonstrate circadian phase-dependent necrotic cell death responses to *Clostridium difficile* toxin B (TcdB). Intriguingly, mouse and human enteroids demonstrate anti-phasic necrotic cell death responses. RNA-Seq data show ~4% of genes are rhythmically expressed in HIEs. Remarkably, we observe anti-phasic gene expression of *Rac1*, a small GTPase directly inactivated by TcdB, between mouse and human enteroids. Importantly, the observed circadian time-dependent necrotic cell death response is abolished in both mouse enteroids and human intestinal organoids (HIOs) lacking robust circadian rhythms. Our findings uncover robust functions of circadian rhythms regulating critical clock-controlled genes (CCGs) in human enteroids governing organism-specific, circadian phase-dependent necrotic cell death responses. Our data highlight unique differences between mouse and human enteroids, and lay a foundation for human organ- and disease-specific investigation of clock functions using human organoids for translational applications.

## Introduction

3-dimensional (3D) organoids are multicellular *in vitro* model systems possessing *in vivo* tissue architecture and function that is unattainable in homogenous single cell culture systems. Multiple tissues have been generated as organoids from a range of sources including pluripotent stem cells (PSC) and patient biopsies^1^. The source utilized for organoid generation influences the maturity of the end sample. For example, PSCs are differentiated through distinct developmental stages, definitive endoderm (DE) and midgut tube formation (MG), to generate human intestinal organoids (HIOs) (**Figure 1A, 1B**); a process mimicking the blastocyst, gastrula, somite and fetal stages of intestinal development, respectively^2,3^. The fetal nature of HIOs is highlighted by the absence of a full spectrum of markers for mature epithelial cell types^4,5^. HIOs can be matured beyond their fetal stage by surgical implantation into the kidney capsule of immunocompromised non-obese diabetic (NOD) *scid* gamma (NSG) mice. When transplanted, they become vascularized by the host circulatory system and develop a well-defined crypt-villus architecture^5^. Transcriptome-wide hierarchical clustering show that transplanted samples are matured to an intermediate state between HIOs and adult intestinal tissue, containing a wider range of intestinal cell type markers compared to HIOs but not equivalent to adult intestinal tissue^4^. Intestinal crypts from the transplanted human tissue can be isolated and cultured *in vitro* to generate mouse kidney capsule-matured human intestinal enteroids (kcHIEs) (**Figure 1A**). Analogously, crypts from human intestinal biopsies can be isolated and grown *in vitro* to generate biopsy-derived HIEs (bHIEs)^6^ (**Figure 1A**). Recent progress highlights the versatile utility of the organoid and enteroid model systems to investigate numerous complex biological questions including the molecular mechanisms maintaining the intestinal stem cell niche^7^, host-pathogen interactions^8^, cellular development^9^ and non-pathogenic bacteria interactions^10^. However, the development and physiological functions of circadian rhythms in human intestinal organoids/enteroids have not been investigated. In this report, we investigated the development of circadian rhythms from PSCs to kcHIEs, and bHIEs to characterize clock-controlled genes (CCGs) and circadian functions in the human intestinal epithelium.

**Figure 1.**
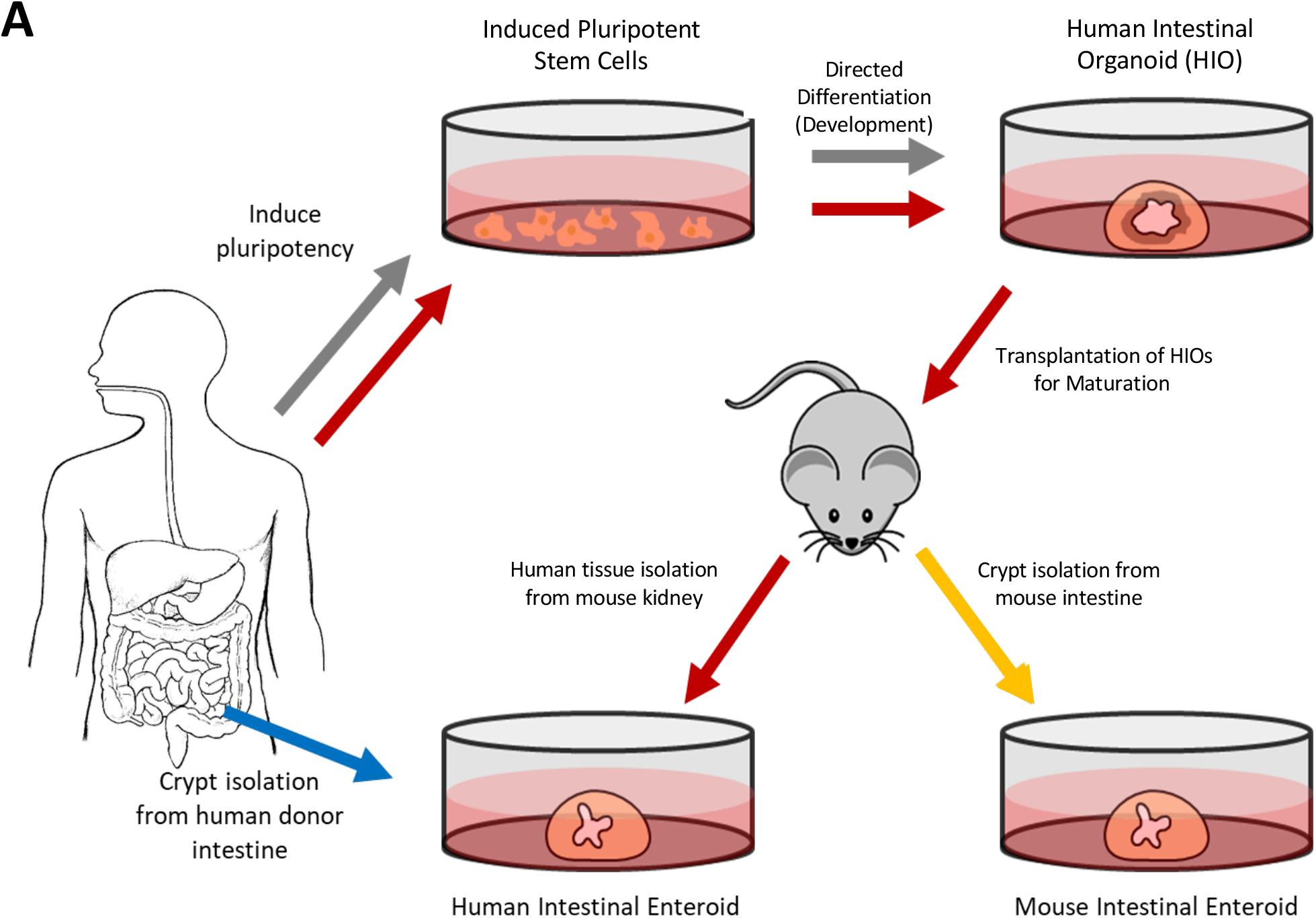
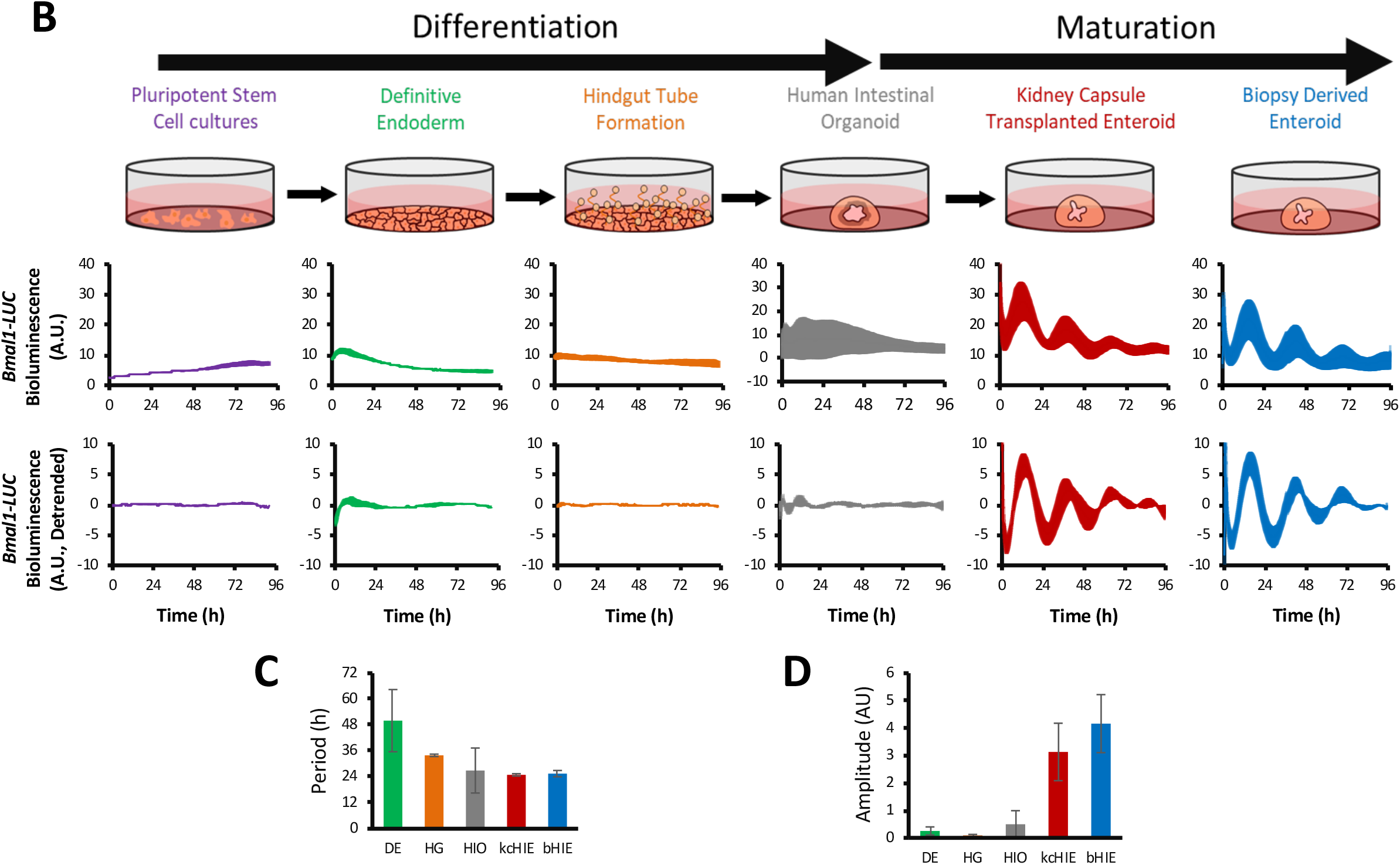
Maturation of HIO is required for the development of robust circadian rhythms. A) Schematic summary for establishing 3D *in vitro* human intestinal organoids (HIOs - grey), kidney capsule matured human intestinal enteroids (kcHIEs - red), biopsy derived human intestinal enteroids (bHIEs - blue) or mouse enteroids (yellow). B) Raw (top) and detrended (bottom) *Bmal1-luc* bioluminescence data throughout directed differentiation of PSCs to HIOs, and kcHIEs and bHIEs. Traces shown are standard deviation of *Bmal1-luc* signal from n=3 biological replicates. C) Period length decreases with continual differentiation of PSCs into HIOs. Maturation of HIOs into kcHIEs leads to 24-hour rhythms with greater reproducibility. D) The amplitude of oscillations are significantly higher in kcHIEs/bHIEs compared to HIOs. *Bmal1-luc* traces in B were generated by plotting the standard deviation of n≥3 biological replicates. C and D depict ± S.D. of n?3 biological replicates. AU stands for arbitrary units.

Circadian rhythms exist throughout the body, providing temporal information to biological processes ranging from metabolism to cell proliferation. Molecularly, circadian rhythms are governed by a transcriptional-translational feedback loop (TTFL), which generates robust circadian oscillations at the single cell level^11^. Two basic helix-loop-helix (bHLH) PER-ARNT-SIM (PAS) domain containing proteins, CLOCK (circadian locomotor output cycles kaput) and BMAL1 (brain and muscle Arnt-like protein-1), form a heterodimeric circadian transcription factor that recognizes E-box motifs on target gene promoters including *Period* (*Per1, Per2, Per3*) and *Cryptochrome* (*Cry1, Cry2*). PER and CRY bind together in the cytoplasm^12^ and translocate into the nucleus where they interact with the CLOCK-BMAL1 heterodimer to repress its activity^13^. PER and CRY are then ubiquitinated and degraded, relieving CLOCK-BMAL1 from negative feedback to re-initiate the cycle^14–16^. CLOCK-BMAL1 forms additional feedback loops with nuclear orphan receptor transcription factors. CLOCK-BMAL1 drive rhythmic expression of positive (*Rorα, Rorβ, Rorγ*) and negative (*Rev-erbα, Rev-erbβ*) nuclear receptors that feedback on *Bmal1* by competitively binding at ROR response elements (RORE) on the *Bmal1* promoter^17–19^.

The suprachiasmatic nucleus (SCN) is commonly referred to as the ‘master clock’ of the mammalian system. Located in the hypothalamus, the SCN contains approximately 20,000 neurons that process photic signals to coordinate circadian rhythms in peripheral organs to external light/dark signals^20,21^. However, peripheral clocks are malleable and can be dissociated from SCN coordination by non-photic circadian zeitgebers. For example, mice restricted to food during their inactive period (i.e. light phase or subjective day) realign the phase of clock and metabolic genes in the liver, but not SCN, to coincide with food availability^22^. Intriguingly, the intestinal microbiota changes over a circadian cycle^23^, and perturbation of the microbiome may impact the function of peripheral circadian rhythms. Antibiotic-induced depletion of microbiota induces constitutively high expression of *Rev-erbα* that abolishes rhythmic toll-like receptor (TLR) expression^24^. Loss of microbiota was also shown to change rhythmic gene expression within the mouse intestinal transcriptome while leaving behavioral rhythms intact^25^. Notably, the recent development of tissue specific conditional *Bmal1* rescue mice was instrumental in uncovering intact peripheral circadian rhythms in the skin epidermis and liver in the absence of circadian signaling from the SCN^26^. These data highlight the importance of peripheral clocks and their ability to process input signals from local factors. More work is needed to uncover fundamental understanding of the conditions and mechanisms that integrate and dissociate the master and peripheral clocks. In this report, we utilized human intestinal organoids and enteroids to characterize the robustness of circadian rhythms during the development of intestinal tissue and investigated the functional role of the intestinal circadian clock in response to *Clostridium difficile* toxin B (TcdB) highlighting a critical function of the endogenous intestinal circadian clock.

*Clostridium difficile* is a spore forming, gram positive, toxin producing intestinal pathogen. *C. diff*. infection (CDI) manifests with diarrhea symptoms that can progress into pseudomembranous colitis and toxic megacolon^27,28^. CDI pathogenesis includes the luminal release of two toxins, toxin A (TcdA) and B (TcdB), that bind to intestinal cellular receptors^29^ and enter the cell via endocytosis^30^. In the cytoplasm, autocatalytic cleavage releases the catalytic glucosyltransferase domain of the toxins^31^ that inactivate the small GTPases RHOA, RAC1, and CDC42, resulting in cellular cytoskeleton breakdown, disruption of epithelial barrier function, and cell death^32,33^. In addition, the NOX1 NADPH oxidase complex has also been implicated in promoting toxin-mediated cell death at high concentrations of TcdB^34^. Several experimental models have been utilized to investigate *C. difficile* pathogenesis and each present limitation. *In vivo* rodent studies have been associated with low reproducibility attributed to confounding variables such as different *C. diff*. strains, antibiotic regimen to induce susceptibility, and cross contamination of animal housing^35^. The translatability of pathophysiological findings from rodent models to humans also cannot be guaranteed. *In vitro* and *ex vivo* studies are, comparatively, controllable but lack multicellular complexity or multi-time point scalability, respectively. Human intestinal organoids/enteroids offer an opportunity to overcome these limitations by coupling *in vitro* experimental control with recapitulation of an *in vivo* human intestinal epithelium. 3D intestinal cultures have been successfully utilized to gain insights into the pathogenic response to human rotavirus^36^, *E. coli^37^* and *Clostridium difficile* toxins A/B^38^. These studies, however, did not investigate the influence of the circadian clock in the pathogenic response. Circadian clock-mediated time of day dependency in the pathogenesis of *Salmonella* Typhimurium^39^ and herpes simplex virus^40^ has been observed in mice, but it is unknown whether humans show identical circadian time-dependent responses via conserved molecular mechanisms. In this report, we utilized mouse and human intestinal enteroids to uncover potential differences with respect to circadian phase-dependent necrotic cell death responses to TcdB.

## Results

### The development of a robust intestinal circadian clock requires maturation of human intestinal organoids

Mouse embryonic stem cells (ESC) do not possess circadian rhythms and have low expression of several canonical clock genes in comparison to NIH3T3 cells^41^. Retinoic acid-induced differentiation of mouse embryonic stem cells initiates autonomous circadian oscillations while reverting differentiated cells back to an induced pluripotent stem cell (iPSC) state abolishes circadian rhythms^42^. Subsequent studies revealed that during the early stages of cellular differentiation, PER1/PER2 are sequestered in the cytoplasm^43^ and the expression of CLOCK is downregulated by *Dicer*/*Dgcr8*-dependent post-transcriptional regulation of *Clock* mRNA^44^ disrupting autonomous oscillations of circadian rhythms. Identical to mouse ESCs and iPSCs, human ESCs and iPSCs lack circadian rhythms and show post-transcriptional suppression of CLOCK protein expression^45,46^. Terminally differentiated cardiomyocytes from human ESCs demonstrate rhythmic gene regulation of clock-controlled genes (CCGs), which regulate circadian time-dependent responses to doxorubicin^45^. Based on these data, we hypothesized that developmental maturation is a critical process for the development of robust circadian rhythms in 3D multicellular intestinal organoids.

To investigate the development of circadian rhythms in multicellular organoids, activity of circadian rhythms was assessed at different stages of development of human intestinal organoids and enteroids using *Bmal1-luciferase* (*Bmal1-luc*) reporter (**Figure 1B**). iPSCs were stably transduced with a plasmid containing *Bmal1-luc* using lentivirus^47^ and differentiated through two distinct stages: definitive endoderm (DE) and midgut tube (MG) formation^2^. *Bmal1-luc* reporter activity was recorded throughout differentiation with a luminometer (KronosDIO^™^, ATTO). In agreement with previous reports^46^, iPSCs did not show circadian oscillations (**Figure 1B**, purple). Interestingly, we observe low intensity bioluminescence signals with long period lengths of about 50 and 34 hours at the DE and MG stages, respectively (**Figure 1B**, **Figure S1**, green and orange). On the other hand, HIOs that were stably transduced with *Bmal1-luc* plasmids demonstrated circadian oscillations with low bioluminescence output (**Figure 1B**, grey). These data show that differentiation of iPSCs into multicellular HIOs promotes period length shortening, culminating with 24-hour bioluminescent oscillations in HIOs (**Figure 1C, 1D**). However, circadian oscillations in these HIOs are not robust, lacking phase uniformity between independent experiments and unable to maintain oscillations longer than 48 hours post-synchronization of circadian rhythms (**Figure 1B, Figure S1**). Although HIOs contain multiple intestinal epithelial cell types^2^, they do not possess the full spectrum of epithelial cell markers, which reflect their immature, fetal-like state. The stem and Paneth cell markers, *Lgr5* and *Lysozyme*, respectively, are expressed in both HIOs and human tissue^4^. The expression of secondary stem cell (*Olfm4*) and Paneth cell (*Defa5, Reg3a*) markers are, however, attenuated in HIOs, indicating they do not possess matured intestinal epithelial cell types^4^. Therefore, we transplanted HIOs into the kidney capsule of NSG mice and allowed them to mature over 3-months^5^ to test if maturation of HIOs could enhance the robustness of circadian rhythms.

After 3-months of maturation, human intestinal crypts were isolated from the grafts to establish *in vitro* mouse kidney capsule-matured human intestinal enteroids (kcHIEs)^6^ (**Figure 1A**). In addition, we generated biopsy-derived human intestinal enteroids (bHIEs) to compare circadian rhythms from matured human samples with HIOs and kcHIEs (**Figure 1A**). Both kcHIEs and bHIEs were transduced with the *Bmal1-luc* plasmid and tested for rhythmic bioluminescent output. In contrast to HIOs, kcHIEs and bHIEs demonstrated robust circadian oscillations, demonstrated by sustained high amplitude oscillations of bioluminescent output from *Bmal1-luc* reporter (**Figure 1B**). To validate the above bioluminescent data, we performed time course experiments collecting HIOs, kcHIEs, and bHIEs every 4-hours over two circadian cycles for real-time quantitative reverse transcription polymerase chain reaction (qRT-PCR). The expression of *Bmal1, Rev-erbα*, and *Per2* was significantly reduced and arrhythmic in HIOs (**Figure 2A**). In contrast, both kcHIEs and bHIEs demonstrated circadian rhythms with appropriate phase relationships between positive (*Bmal1*) and negative (*Per2, Rev-erbα*) clock elements (**Figure 2B, 2C**). Interestingly, the average amplitude of *Bmal1, Per2* and *Rev-erbα* are significantly lower in kcHIEs compared to bHIEs (**Figure 2D**), which suggest that bHIEs possess a more robust circadian clock compared to kcHIEs. The average expression of these genes are not significantly different between kcHIEs and bHIEs except for *Rev-erbα* (**Figure 2D**). Based on these data, we hypothesize that circadian time-dependent responses to external perturbations will be observed in kcHIEs and bHIEs, but not in HIOs. As a proof of principle, we tested the circadian clock-dependent response to *Clostridium difficile toxin B* (TcdB).

**Figure 2.**
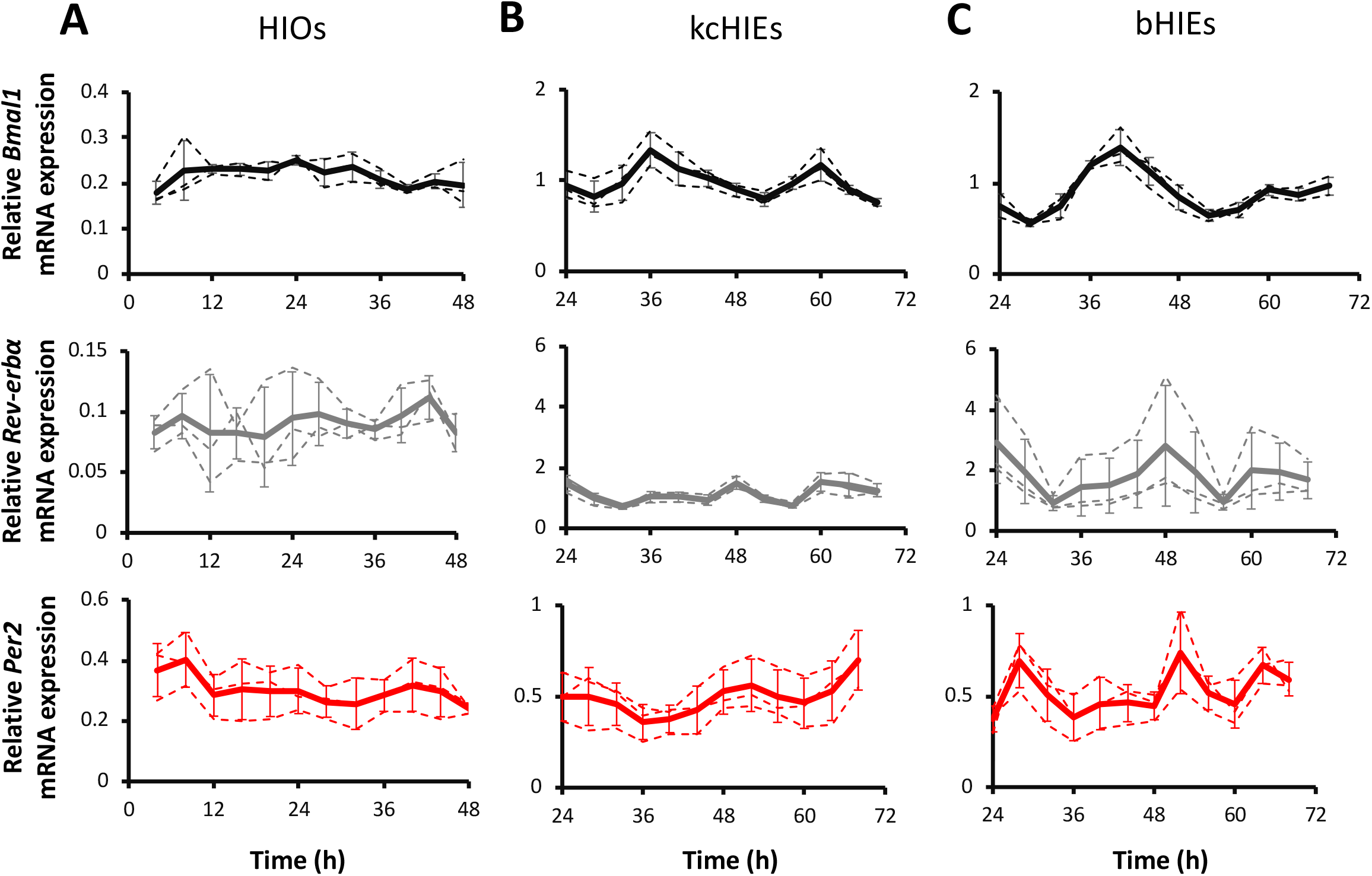
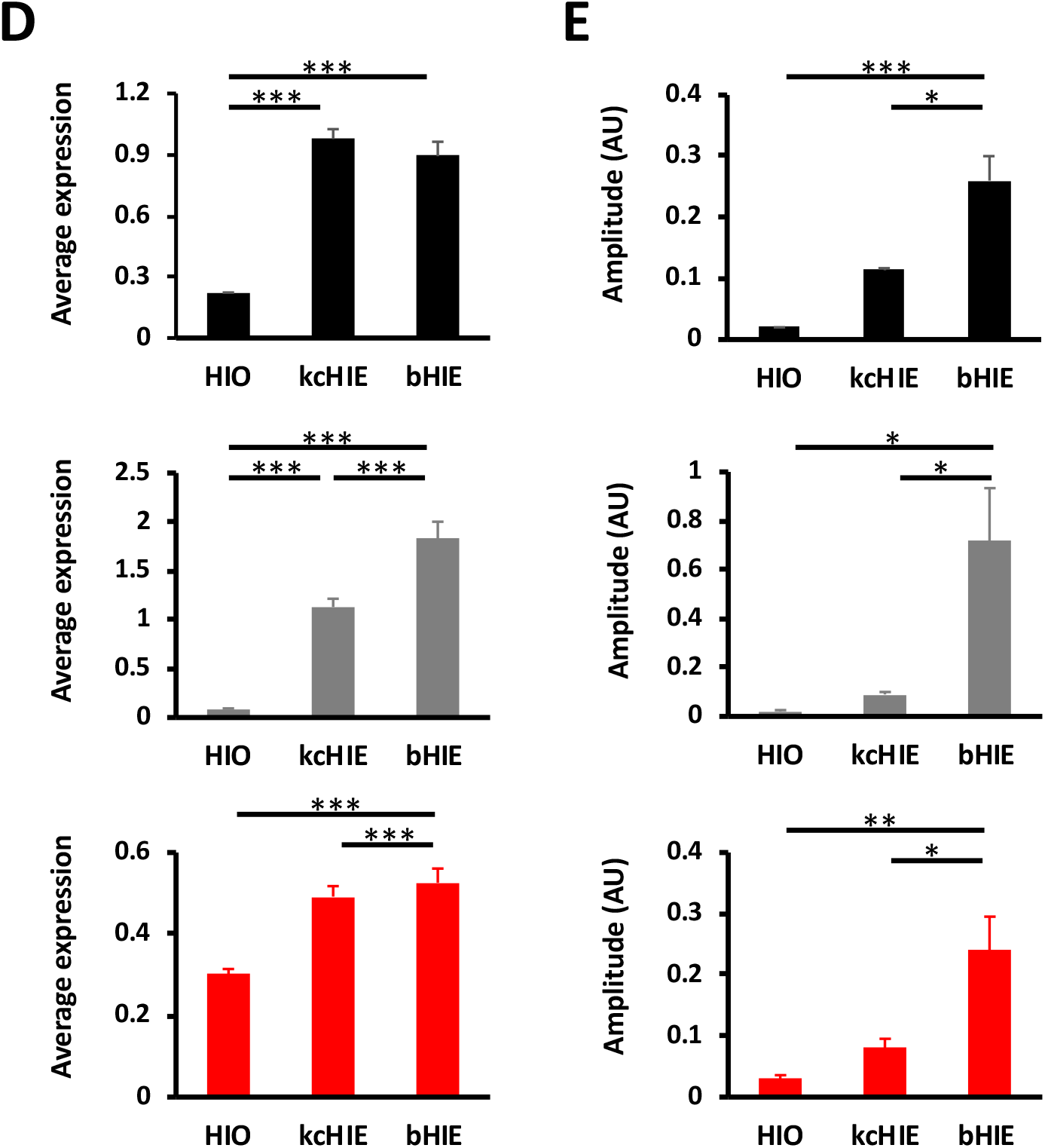
kcHIEs and bHIEs show circadian rhythms with appropriate phase relationships between canonical clock genes. A-C. mRNA expression of *Bmal1* (black), *Rev-erbα* (grey) and *Per2* (red) in HIOs (A), kcHIEs (B) and bHIEs (C). D. Average expression of *Bmal1* (black), *Rev-erbα* (grey) and *Per2* (red) in HIO, kcHIE, and bHIE. E. Amplitude of *Bmal1* (black), *Rev-erbα* (grey) and *Per2* (red). AU stands for arbitrary units. The data in A-C are shown as mean (bold line) ± S.D. with replicates displayed as dashed lines, and the data in D-E are shown with ± S.D. of n=3 biological replicates.

### The intestinal circadian clock in mouse and human enteroids regulate circadian phase-dependent necrotic cell death responses to TcdB

Circadian clock-dependent time-of-day variation in the pathogenic response has been demonstrated for *Salmonella enterica serovar* Typhimurium^39^, herpes and influenza A virus^40^ and *Leishmania*, in mice^48^. However, circadian clock-dependent responses to bacterial toxins in mouse or human enteroids have not been tested. It is critical to determine circadian clock-dependent responses to different external perturbations (e.g. pathogens, toxins, nutrients, etc.) in both mouse and human enteroids to compare and contrast their responses in order to develop potential translational applications using patient-derived HIEs while taking advantage of various tools in transgenic mice. Therefore, we assessed circadian clock-dependent responses following exposure to purified TcdB in both mouse and human enteroids.

Mouse enteroids were isolated from the tamoxifen-inducible *Bmal1^f/f^-EsrCRE* mouse^49^. *In vitro* treatment of tamoxifen in *Bmal1^f/f^-EsrCRE* enteroids successfully abolished circadian rhythms (**Figure S2**), which enabled us to use *Bmal1-floxed* enteroids as arrhythmic negative controls. Circadian rhythms of *Bmal1^f/f^-EsrCRE* enteroids were synchronized using dexamethasone, and 10ng/mL of TcdB was added to the growth media at 24- (D24) and 36-hour (D36) post-dexamethasone treatment (**Figure S3**). SYTOX^™^ Orange, a fluorescent dye that stains nucleic acids in cells with disrupted membranes, was added simultaneously with TcdB to quantify necrotic cell death at 2, 24, and 48 hours post-exposure of TcdB. In control *Bmal1^f/f^-EsrCRE* enteroids, without tamoxifen treatments, we observe greater necrotic cell death in the D36 group compared to D24 at 48 hours post TcdB treatments (**Figure 3A**). Identical data was observed in mouse enteroids derived from PER2::LUC mice (**Figure S3**). In contrast, tamoxifen-treated *Bmal1-floxed* enteroids showed a loss of circadian time-dependent necrotic cell death responses to TcdB (**Figure 3B, 3D**), which indicates that the intestinal circadian clock regulates the observed circadian phase-dependent phenotypes. Next, we used the aforementioned three human organoid models to test whether this phenotype from mouse enteroids is conserved in human organoids.

**Figure 3.**
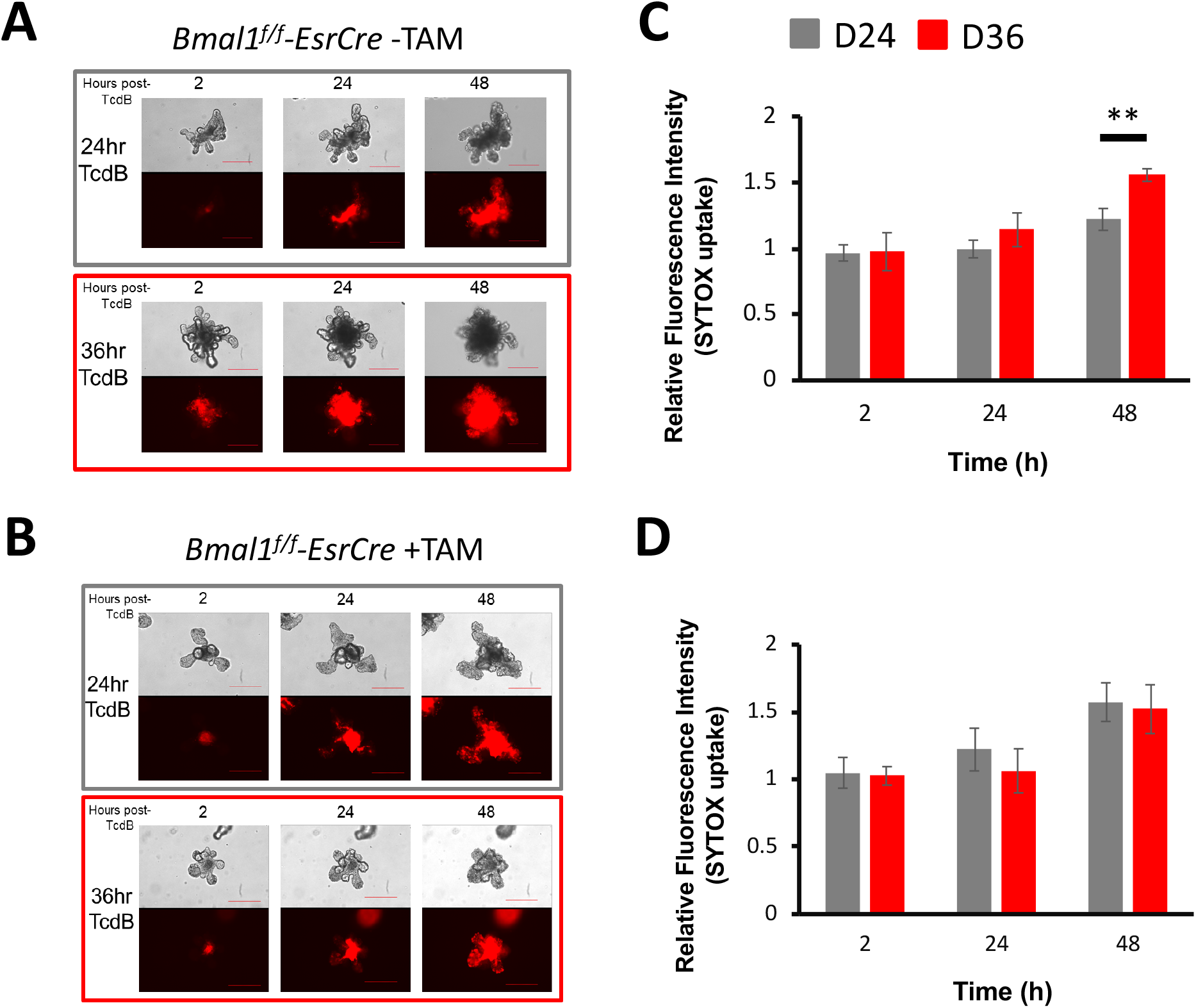
The mouse enteroids show circadian clock-dependent necrotic cell death response to *Clostridium difficile* toxin B. A-B) Representative images of necrotic cell death (SYTOX orange - red fluorescence) at 2-, 24- and 48-hours post exposure to 10ng/mL TcdB in control *Bmal1^f/f^-EsrCRE* without tamoxifen (A) or arrhythmic *Bmal1^f/f^-EsrCRE* with tamoxifen-treated (B) enteroids. Mouse enteroids were exposed to TcdB at two circadian timepoints, 24-hours (D24, grey) or 36-hours (D36, red) post-synchronization. C-D) Quantitative analysis of fluorescent intensity from SYTOX orange in control *Bmal1^f/f^-EsrCRE* without tamoxifen (C) or arrhythmic *Bmal1 ^f/f^-EsrCRE* with tamoxifen-treated (D) enteroids. Data represented as mean ± S.D. of n=4 biological replicates (i.e. enteroids derived from different mice). All of the data were normalized to time and phase matched vehicle controls set to 1. **p<0.01. Scale bar = 250μM.

HIOs, kcHIEs and bHIEs were tested for circadian clock phase-dependent responses following the same protocol used for mouse enteroids. HIOs showed identical necrotic cell death responses between the D24 and D36 groups (**Figure 4A**) consistent with their lack of robust circadian rhythms (**Figure 1**). In contrast, both kcHIEs and bHIEs demonstrated circadian phase-dependent necrotic cell death responses to TcdB with greater amount of cell death in the D24 compared to the D36 group for both kcHIEs and bHIEs (**Figure 4B**). These findings indicate that a functional, robust circadian clock is required for circadian phase-dependent responses to TcdB. Intriguingly, mouse and human enteroids demonstrated anti-phasic necrotic cell death responses to TcdB. Mouse enteroids showed a higher necrotic cell death in the D36 group (**Figure 3B**) whereas HIEs were more responsive in the D24 group (**Figure 4E, 4F**). We speculated that anti-phasic behavioral rhythms between diurnal humans and nocturnal mice could be hardwired within the peripheral circadian clocks determining distinct phases of CCGs between mouse and human enteroids. To uncover CCGs in HIEs and identify potential target genes for the observed anti-phasic necrotic cell death responses in mouse vs. human enteroids, we performed a 48-hour time course experiment with 2-hour resolution extracting RNA from bHIEs for RNA-Seq.

**Figure 4.**
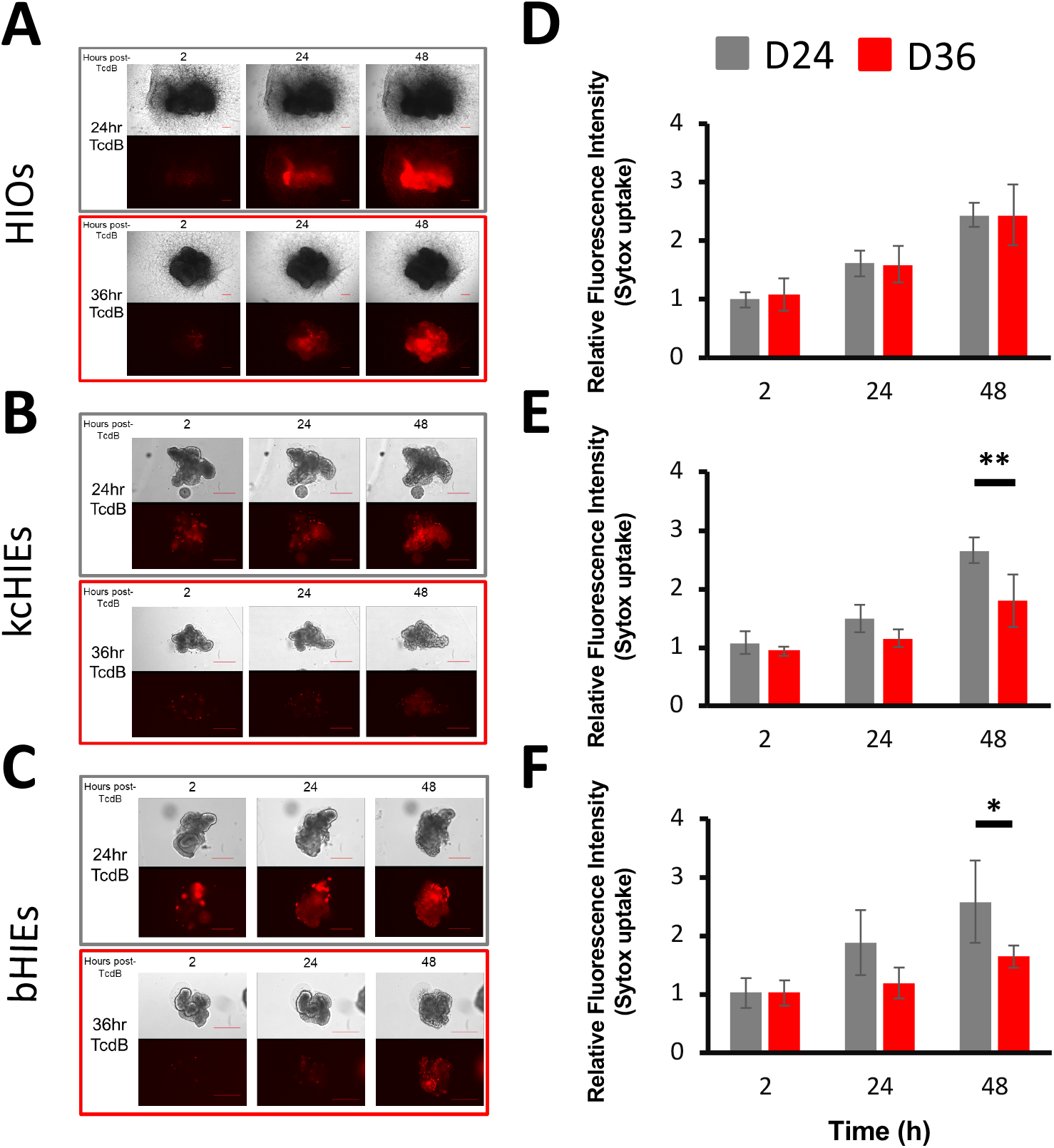
A functional circadian clock is needed for a circadian-dependent response to TcdB in human organoids. A-C) Representative images of necrotic cell death (SYTOX orange) at 2-, 24- and 48-hours post exposure to 0.1μg/mL, 0.5μg/mL and 0.5μg/mL TcdB in HIOs (A), kcHIEs (B) and bHIEs (C), respectively. D-F) Quantification of necrotic cell death in HIOs (D), kcHIEs (E) and bHIEs (F). All samples were exposed to TcdB at two circadian timepoints, 24-hours (D24, grey) or 36-hours (D36, red) post-synchronization. Data are shown as mean ± S.D. of n?3 biological replicates (i.e. HIOs are generated from different PSCs and bHIEs are derived from three different patients’ samples). All data were normalized to time and phase matched vehicle controls set to 1. *p<0.05, **p<0.01. Scale bar = 250μM.

### Approximately 4% of genes in human intestinal enteroids show rhythmic gene expression

For the analysis of our HIE RNA-Seq data, we used MetaCycle^50^ to identify clock controlled genes (CCGs) and performed phase set enrichment analysis (PSEA)^51^ to determine signaling pathways that are under the control of intestinal circadian clocks. For MetaCycle analysis, we excluded low expression genes with mean expression below the lowest 25%, and used p-value cutoff with base expression >0.05 and relative amplitude > 0.01. These analyses uncovered 730 CCGs out of 18,039 genes, which is approximately 4.05% of total genes above the cutoff, with a highest number of genes peaking during the early phase post-clock synchronization (**Figure 5A, B**). This is in sharp contrast to less than a dozen cycling genes in other *in vitro* systems (i.e. NIH3T3 and U2OS cells)^52^. Importantly, heatmap of Spearman’s rho for circadian clock and clock-associated genes show appropriate correlation of each gene against the rest (**Figure 5C**), which confirmed that the circadian clock machinery is intact and functional with proper phase relationships in HIEs.

**Figure 5.**
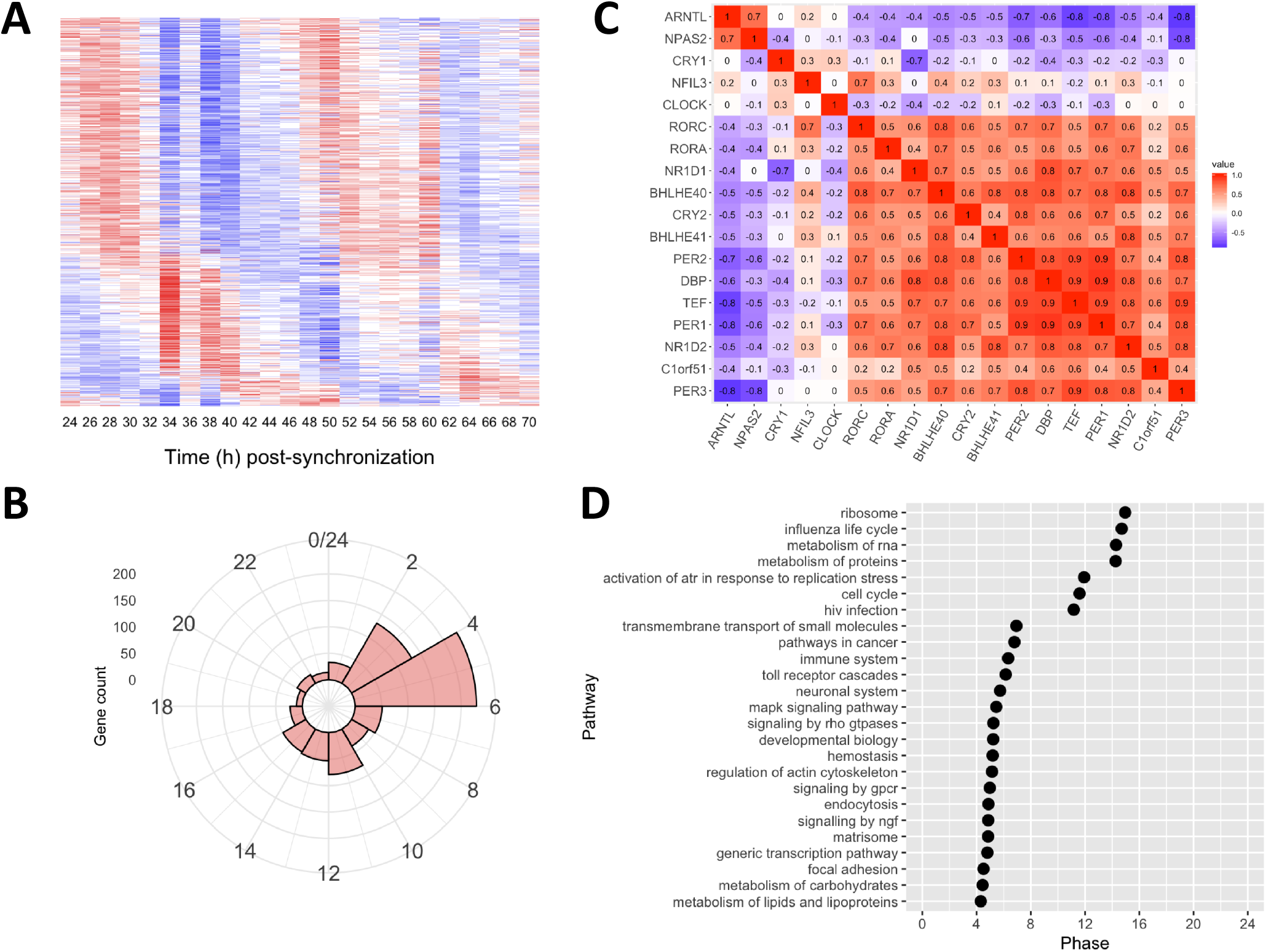
Approximately 4% of genes show circadian oscillations in HIEs. A) Heatmap of circadian genes over 48 hours in HIEs. B) Phase plot demonstrates the number of rhythmic genes across different phases of circadian rhythms. C) Spearman’s rho for circadian clock and clock-associated genes. D) 25 signaling pathways identified by phase set enrichment analysis (PSEA).

PSEA uses prior knowledge to identify signaling pathways demonstrating temporally coordinated expression profiles. PSEA revealed that the intestinal epithelial circadian clocks regulate temporal organization of 25 signaling pathways with three distinct clusters of genes peaking at different circadian phases, which are associated with metabolism, cell cycle, immune system, and cancer (**Figure 5D**). Interestingly, the genes that have a major role in regulating the development and homeostasis of intestine, such as *FoxA4, Wnt3*, and *Bmp4*, were observed in the early phase. Importantly, PSEA uncovered 10 genes that are involved in signaling by RHO GTPases, which included a single TcdB target gene, *Rac1*. Other TcdB target genes, *Cdc42* and *RhoA*, did not show rhythmic gene expression profiles in HIEs (**Figure 6A, B**). To our surprise, we observe anti-phasic expression of *Rac1* in human and mouse enteroids RNA-Seq data (**Figure 6C**). Hence, we validated the rhythmic expression of *Rac1* in HIEs and compared its expression against mouse enteroids using qRT-PCR. Consistent with our RNA-Seq data, we observe anti-phasic expression of *Rac1* in mouse and human enteroids (**Figure 6D**). Lastly, we observe arrhythmic expression of *Rac1* in both HIOs and *Bmal1-floxed* mouse enteroids, which is consistent with lack of circadian phase-dependent necrotic cell death responses (**Figure S4**). Together, our data strongly suggest that the rhythmic expression of *Rac1* determines the observed anti-phasic necrotic cell death response against TcdB in mouse vs. human enteroids.

**Figure 6.**
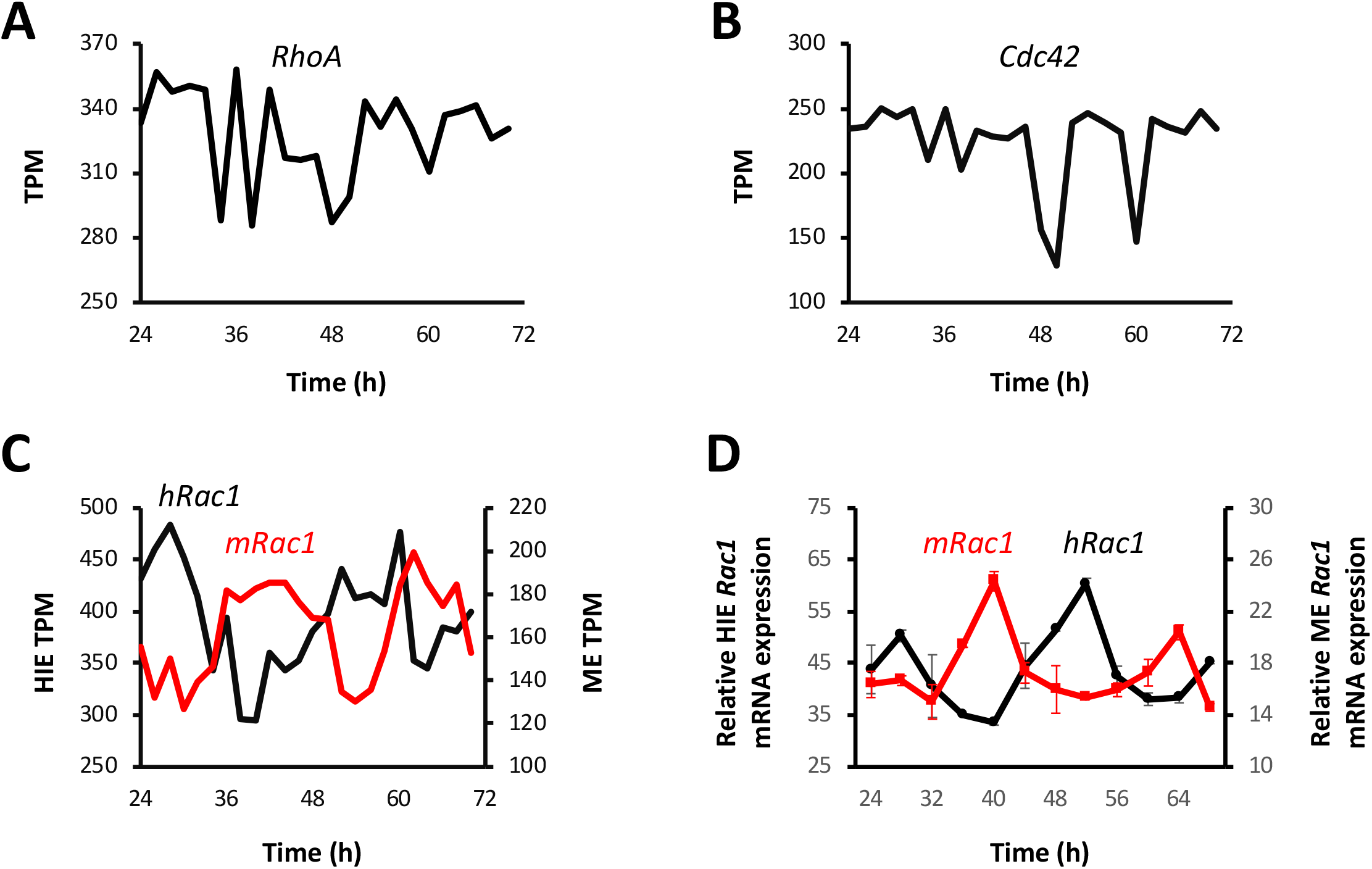
*Rac1* gene expression shows anti-phasic profiles between mouse and human enteroids. A-B) TcdB target small GTPases, *RhoA* and *Cdc42*, do not show circadian oscillations in HIEs. C) *Rac1* expression shows anti-phasic profiles in mouse (*mRac1*, red) and human enteroids (*hRac1*, black) RNA-Seq data. D) Validation of anti-phasic *Rac1* expression in mouse (*mRac1*, red) and human enteroids (*hRac1*, black) by qRT-PCR. TPM stands for transcripts per million.

## Discussion

Circadian rhythms regulate CCGs in a tissue-specific manner throughout the body^53^. Circadian rhythms in peripheral organs are affected by numerous inputs including light^54,55^, nutrients^22,56,57^, and systemic cues from SCN^58–62^. Previous studies characterizing CCGs in different organs from *in vivo* mice uncovered organ-specific regulation of circadian rhythms integrated by both peripheral clocks and systemic cues from SCN. Recent development of tissue-specific conditional *Bmal1* rescue mice will be instrumental to discover organ-specific endogenous CCGs independent of SCN. Characterization of organ-specific endogenous CCGs is necessary to systematically compare and contrast changes of circadian gene expression profiles in the presence of systemic and external cues, and control and disease states (e.g. cancer). However, the discovery of organ-specific endogenous CCGs in humans is difficult, creating a major bottleneck hindering organ-and disease-specific investigation of circadian rhythms and CCGs for translational applications.

Human organoids are complex, multicellular *in vitro* systems that are devoid of systemic signaling factors, making them an appealing system to evaluate endogenous, tissue-specific functions of peripheral circadian rhythms. The function of circadian rhythms, however, will depend on the maturity of organoids, because previous studies indicated lack of circadian rhythms in PSCs and mouse fetus during the early stages of its development^45,46,63^. Therefore, we first characterized robustness of circadian rhythms from iPSCs to HIOs demonstrating the lack of robust circadian rhythms in HIOs. Differentiation of mouse embryonic or induced pluripotent stem cells results in the onset of circadian oscillations after 28 days^44^. Interestingly, a subsequent study from the same group found that human iPSCs require up to 90 days of differentiation to initiate circadian oscillations^46^. On the other hand, differentiation of human embryonic stem cells (ESC) into cardiomyocytes results in rhythmic cultures oscillations after 30 days. However, it is worth indicating that circadian oscillations of *Bmal1-dLuc* and *Per2-dLuc* reporter activities in the differentiated cardiomyocytes show weak oscillations^45^ compared to HIEs (**Figure 1**) and differentiated cells from iPSCs after 90 days of differentiation^46^. The lack of robust circadian rhythms in HIOs after 35-day differentiation protocol is consistent with the lack of circadian oscillations at day 42 of *in vitro* differentiation of human iPSCs by Umemura and colleagues^46^, which indicate that further maturation is necessary for the development of circadian rhythms.

In contrast to HIOs, we observe robust circadian rhythms in both kcHIEs and bHIEs based on their identical bioluminescence outputs and oscillations reflecting the activity of *Bmal1-luc* reporter (**Figure 1**). However, we observe lower amplitude of *Bmal1, Per2* and *Rev-erbα* transcripts in kcHIEs compared to bHIEs (**Figure 2D**). These findings suggest that kcHIEs represent an intermediate state between HIOs and bHIEs, which is in agreement with previous results^4,5^. Transcriptomic analysis comparing HIOs, kcHIEs, and adult intestinal tissue demonstrated that kcHIEs possess intermediate expression of brush border enzymes, *sucrase isomaltase* and *trehalase*, between HIOs and adult intestinal tissue^5^. Similar findings were discovered for expression profiles of Paneth and stem cell markers in HIOs, kcHIEs, vs. adult intestinal tissue^4^. It would be interesting to perform RNA-Seq using time course samples from kcHIEs to compare and contrast CCGs between kcHIEs and bHIEs characterizing the reorganization of CCGs during development.

Approximately 4% of transcriptome show circadian oscillations in bHIEs regulating diverse signaling pathways ranging from cell cycle to immune response. The number of cycling genes in HIEs is similar to the oscillating transcripts in human embryonic stem cell-derived cardiomyocytes^45^, which indicate that these *in vitro* systems possess functional circadian rhythms regulating a large number of tissue-specific CCGs. In contrast, NIH3T3 and U2OS cells show less than a dozen cycling genes despite the presence of robust core clock gene oscillations^52^. On the other hand, 3 – 16% of cycling genes are observed depending on peripheral organs of mice *in vivo^64^*. We hypothesize that intestinal epithelial cells *in vivo* will show a greater number of CCGs compared to enteroids, because those CCGs will be a result of both systemic cues and peripheral circadian clocks. In our future work, we plan to compare CCGs in mouse enteroids and mouse intestinal tissue to systematically dissect CCGs that are controlled by systemic cues and intestinal clocks.

The intestinal circadian clock possesses robust circadian rhythms^65^, and is responsible for promoting circadian variation in a wide range of biological processes^66^, including intestinal motility^67^, nutrient absorption^68^, innate immunity^69^, timing of cell division^70^, and the pathogenic response^39^. In patients, nocturnal diarrhea is an “alarm” symptom that should prompt clinicians to consider further evaluation, including testing for *C. difficile* infection^71^. However, whether a disrupted circadian clock predisposes to or is a result of gastrointestinal infections remains largely unknown. Insights to such questions will be needed to determine whether or not targeting circadian rhythms to treat such diseases would be efficacious. Our findings demonstrate robust operation of canonical clock genes regulating specific signaling pathways, and species-dependent anti-phasic expression of *Rac1* between mouse and human enteroids explaining the observed anti-phasic necrotic cell death response in mouse vs. human enteroids against TcdB. In the future, we plan to evaluate the importance of circadian-phase specific expression of *Rac1* and other genes that are involved in signaling by RHO GTPases between mouse and human enteroids, and design chronotherapeutic regimens for treatments of *C. difficile* infection. Furthermore, our work demonstrates a proof of concept that human organoids derived from different organs and disease states could provide a unique opportunity to: 1) characterize function of peripheral circadian clocks in different organs, 2) identify circadian biomarkers for human diseases, and 3) leverage the information to develop novel circadian treatment regimens for translational applications.

## Acknowledgement

We thank support from the Digestive Health Center (NIH P30 DK078392) at CCHMC and the Live Microscopy Core at University of Cincinnati College of Medicine. This work was supported by U19 AI116491 (JMW, SRM, CIH), U19 AI116497 (NFS), R01 DK117005 (CIH), P01 HD093363 (JMW), UG3 DK119982 (JMW), University of Cincinnati Bridge Funding, and CCHMC Research Innovation and Pilot Funding.

A.E.R and M.P. designed the experiments, collected the data, generated and compiled figures and wrote the manuscript. T.M. and S.L. assisted in designing and performing experiments. K.R.S., K.C, and N.S. received and aligned RNA-sequencing data to reference transcriptomes. D.E.F., G.W. and J.B.H. analyzed RNA-sequencing data and generated the associated figures. N.S. and J.A.H. performed surgical transplantation of HIOs into the kidney capsule of mice and provided kcHIEs. T.R.B. provided HIOs and assisted in the design of HIO related experiments. H.A.M. provided kcHIE samples. N.F.S. supplied bHIEs and reviewed the manuscript. M.A.H, J.M.W and J.B.H. were involved in the study design and review of this manuscript. S.R.M. and C.I.H. designed and directed the overall progress of studies, wrote and provided final approval of the manuscript.

## Materials & Methods

### Animals

Mouse enteroids were established and cultured from 3-6 month old C57BL/6J (Jackson Laboratories), PER2::LUCIFERASE^72^ and *Bmal1^f/f^-EsrCRE*^49^. Immune-deficient NSG mice (Jackson Laboratories) were utilized for kidney capsule transplantation of HIOs^5^. Animal handling was performed according to IACUC protocols #2016-0014 (PI: Michael Helmrath - Cincinnati Children’s Hospital Medical Center) and #17-01-30-01 (PI: Christian Hong - University of Cincinnati).

### Isolation and maintenance of mouse enteroids

Mouse jejunal enteroids were generated following established protocols^73^. Briefly, crypt domains from small intestinal tissue were isolated, added to 40μL of Matrigel (Corning) membrane matrix, suspended as a single 3D Matrigel dome within a 24-well plate and provided 400μL mouse enteroid media (Advanced DMEM/F-12 (Gibco), L-Glutamine (Gibco), Penicillin/Streptomycin (Gibco), N-2 (Gibco), B-27 (Gibco), 10mM HEPES solution (Millipore-Sigma), 50ng/mL recombinant murine EGF (Peprotech), and R-Spondin/Noggin conditioned media generated inhouse. Enteroids were passaged every 4-7 days via a 27G syringe (BD). CRE driven *Bmal1* exon 4 excision was initiated *in vitro* by adding 1μM 4-OH tamoxifen (Cayman) for 24-hours to *Bmal1^f/f^-EsrCRE* enteroids. All samples were maintained in cell culture incubators set at 37°C and 5% CO_2_. Enteroids isolated from separate mice were considered biological replicate samples.

### Experiment design to test for a circadian phase-dependent response to TcdB

1-2 HIOs and ≈15-20 mouse/human enteroids were plated per well to control for density. 100nM dexamethasone (DEX) was added to treatment groups either 36-hours (D36) or 24-hours (D24) prior to TcdB addition. Samples were then exposed to a vehicle buffer (50mM Tris Base Ultrapure (US biological), 100mM NaCl (Fisher Scientific), pH 7.5) or *Clostridium difficile* toxin B (List Laboratories) by addition to cell culture media. SYTOX Orange (500nM) (Invitrogen) was added with Vehicle/TcdB for quantification of necrotic cell death. Images were acquired at 2-, 24- and 48-hours post-Vehicle/TcdB addition on a Dmi8 inverted microscope (Leica Microsystems). Quantification of sytox uptake (necrosis) was performed using the LasX software (Leica Microsystems) area of interest feature to quantify fluorescence within individual organoid/enteroids. All TcdB data were set relative to time- and phase-matched vehicle controls set to 1.

### HIO generation

HIOs were generated following previously established protocols^2^ using multiple pluripotent stem cell (PSC) lines, both induced and embryonic. Briefly, PSCs were differentiated through the definitive endoderm stage via the addition of Activin A (Cell Guidance Systems) between day-0 (D0) and D2, BMP4 (R&D Systems) at D1 and CHIR99021 (Stemgent) / FGF4 (R&D Systems) from D3 to D6. The addition of CHIR99021/FGF4 prompted the formation of midgut tube spheroids from the definitive endoderm monolayer. At D7 midgut tube spheroids were collected and suspended in a Matrigel dome (Corning). Spheroids were then grown in gut media: DMEM/F12 (Gibco) supplemented with N-2 (Gibco), B-27 (Gibco), HEPES solution (Millipore-Sigma), recombinant murine epidermal growth factor (Peprotech), L-Glutamine (Gibco), Penicillin/Streptomycin (Gibco). At D21 sample density was reduced to ≈1-3 per 50μL Matrigel bubble. Samples were grown for an additional 14-days in gut media, resulting in terminal HIO cultures^2^. Individual HIOs were considered as biological replicates for all experiments.

### Generation and maintenance of kidney capsule-matured- and biopsy derived-human intestinal enteroids

HIOs were transplanted into the kidney capsule of immune-deficient NOD-SCID IL-2Rγ^null^ (NSG) (Jackson Laboratories) mice, as previously described^5^. Crypt domains from transplanted HIOs were isolated and suspended in Matrigel domes to form kidney capsule-matured human intestinal enteroids (kcHIEs)^6^. Frozen duodenal patient biopsy derived human intestinal enteroids (bHIEs) were provided by the lab of Dr. Noah Shroyer (Baylor College of Medicine). Both bHIEs and kcHIEs were cultured following identical protocols. Enteroids were expanded by suspending pelleted enteroids in Matrigel and plating as three separate 10μL domes/well in a 24-well plate. Intesticult organoid growth medium (StemCell Technologies) was used for enteroid expansion. Human enteroid differentiation was initiated by replacing expansion media with differentiation (Intesticult Component A (StemCell Technologies) mixed at a 1:1 ratio with DMEM/F12 supplemented with 15mM HEPES) media 2-days post-passaging. Human enteroids were maintained in differentiation media at least four days before being used for experiments. kcHIEs derived from separate transplanted mice were counted as biological replicates. bHIEs isolated from separate human patient biopsies were counted as biological replicates.

### Bmal1-luciferase lentiviral transduction and monitoring of Bmal1 activity

All human samples from PSCs to HIOs, kcHIEs and bHIEs were transduced with the same pABpuro-BluF (Addgene) plasmid DNA^47^. Plasmid DNA was packaged into lentiviral vectors by the Viral Vector Core at Cincinnati Children’s Hospital Medical Center (CCHMC). PSC transduction was carried out following a previously published lentiviral transduction protocol^70^. Transduced PSCs were Puromycin selected (2μg/mL) (Invivogen) and differentiated into definitive endoderm (DE) and midgut tube (MG) cultures, generating PSC, DE and MG clock reporting samples. Midgut tube samples were re-transduced following the same protocol used for PSC transduction and differentiated into HIOs to generate *Bmal1-luc* reporting HIOs. The bHIE and kcHIE transduction protocol was designed by merging previously reported enteroid digestion^74^ and transduction^75^ protocols. After transduction, digested enteroids were suspended in Matrigel and plated following normal culture protocol. 350μL of Intesticult Component A/B (1:1) media supplemented with 10μM ROCK inhibitor and 10μM CHIR99021 was added to the culture. After 24-hours, the media was replaced with 350μL Intesticult Component A/B (1:1) supplemented with 10μM ROCK inhibitor. On day two post-infection media was transitioned to Intesticult Component A/B (1:1) supplemented with 2μg/mL Puromycin. Puromycin selection was maintained for 2-weeks before starting experiments. *Bmal1-luc* transduced samples were monitored for clock output over 4-days with a KronosDIO luminometer (ATTO). PSC, DE, MG and HIO samples were maintained in normal culture media throughout recording. kcHIEs and bHIEs were differentiated for 4-days prior to bioluminescence recording. On day 4, samples were re-plated to 35mm dishes, synchronized and added to the KronosDIO to begin recording. All samples were provided a 1-hour clock synchronization stimulus with 100nM DEX (Millipore-Sigma) and given new, non-DEX containing, media prior to recording. 200μM Luciferin (GoldBio) was added to each culture for luminescence detection.

### Time course sample collection

Rhythmic gene expression was determined via time course sample collection. Samples were collected every 4-hours over a 48-hour period, two full circadian cycles. At each collection timepoint, cells were pelleted, suspended in 1mL TRI reagent (Molecular Research Center) and snap frozen in liquid N2. After completion of the time course, RNA was isolated via a RNeasy mini-kit column (Qiagen) and cDNA were generated using the Superscript III Reverse transcriptase kit (Invitrogen). FAST SYBR green (Applied Biosystems) and gene specific primer pairs were used to quantify gene expression via real-time quantitative reverse transcriptase polymerase chain reaction (qRT-PCR) with TATA-Box Binding protein (*Tbp*) expression used as a housekeeping gene.

### RNA-sequencing analysis

To perform RNA-sequencing, we collected C57BL/6J mouse enteroids and human bHIEs as a time course. Mouse enteroids were pooled from two separate mice to mitigate single mouse biasing and bHIEs were from a single patient (female, 24-years old) biopsy. Samples were collected every two hours for 46-hours starting at 24-hours post DEX synchronization. Isolated RNA samples were shipped on dry ice for further processing and sequencing analysis using paired-end reads with 50 million reads per sample by Novogene. Raw RNA-sequencing FASTQ files were directly used to generate expression files using the Kallisto^76^ embedded algorithm inside the Altanalyze module^77^. Kallisto pseudo-aligned each transcript with the Ensemble reference transcriptome (Ensembl version 72) and calculated TPM (transcripts per million) by Altanalyze^77^.

### Statistical analysis

All data are displayed as mean values of at least three biological replicates with error bars representing standard deviation around the mean. KronosDIO bioluminescent traces were quantified for clock parameters via Fast-Fourier Transformation^70^. qRT-PCR and RNA-seq transcript expression profiles were tested for significant rhythms using the R package MetaCycle 2D^50^ (p-value cutoff of 0.05). All between group comparisons were made using a two-way analysis of variance (Two-Way ANOVA) with Tukey’s post-hoc analysis. Normality of the residuals was confirmed with Shapiro’s test of residuals at a significance of p<0.05. All statistical tests were performed in R Studio version 1.2.1335.

