## Supplementary Figures for "Ontogeny and function of the circadian clock in intestinal organoids"

**Figure S1** - Detrended bioluminescent data from pluripotent stem cell (PSC), definitive endoderm (DE), midgut tube spheroids (MG) and HIOs (A, B, C and D, respectively) with zoomed-in Y-axis. AU stands for arbitrary units**.**

**Figure S2** – Tamoxifen-treated *Bmal1$^{f/f}$-EsrCRE* mouse enteroids shows arrhythmic clock gene expression. A-B) Control mouse enteroids, *Bmal1$^{f/f}$-EsrCRE* without tamoxifen treatment, show rhythmic *Bmal1* (black) and *Rev-erbα* (grey) gene expression with out-of-phase profiles.

C-D) *Bmal1* (black) and *Rev-erbα* (grey) gene expression are low and arrhythmic in tamoxifen-treated *Bmal1$^{f/f}$-EsrCRE* mouse enteroids. Thick lines are mean ± S.D of n=3 biological replicates depicted as dotted lines.

**Figure S3** - The circadian phase-dependent necrotic cell death response to TcdB is observed in control PER2::LUC mouse enteroids. A) Schematic summary of the experimental design to test for a circadian phase-dependent response to TcdB *in vitro*. Separate samples were synchronized either 36-hours or 24-hours prior to TcdB exposure to generate two sample groups 12-hours out-of-phase at the time of TcdB addition. B) Representative images of necrotic cell death (SYTOX orange - red fluorescence) at 2-, 24- and 48-hours post exposure to 10ng/mL TcdB in PER2::LUC enteroids. C) Quantitative analysis of fluorescent intensity from SYTOX orange in PER2::LUC mouse enteroids. PER2::LUC enteroids show greater necrotic cell death in the 36-hour group compared to the 24-hour group. Data represented as mean ± S.D. of n=4 biological replicates (i.e. enteroids derived from different mice). All data were normalized to time and phase matched vehicle controls set to 1.**p<0.01. Scale bar = 250μM.

**Figure S4** - *Rac1* gene expression is arrhythmic in HIOs and *Bmal1-floxed* mouse enteroids. *Rac1* gene expression in HIOs (A) and *Bmal1-floxed* mouse enteroids (B). The data are shown as mean (bold line) ± S.D. with replicates displayed as dashed lines with ± S.D. of n=3 biological replicates.

### Figure S1

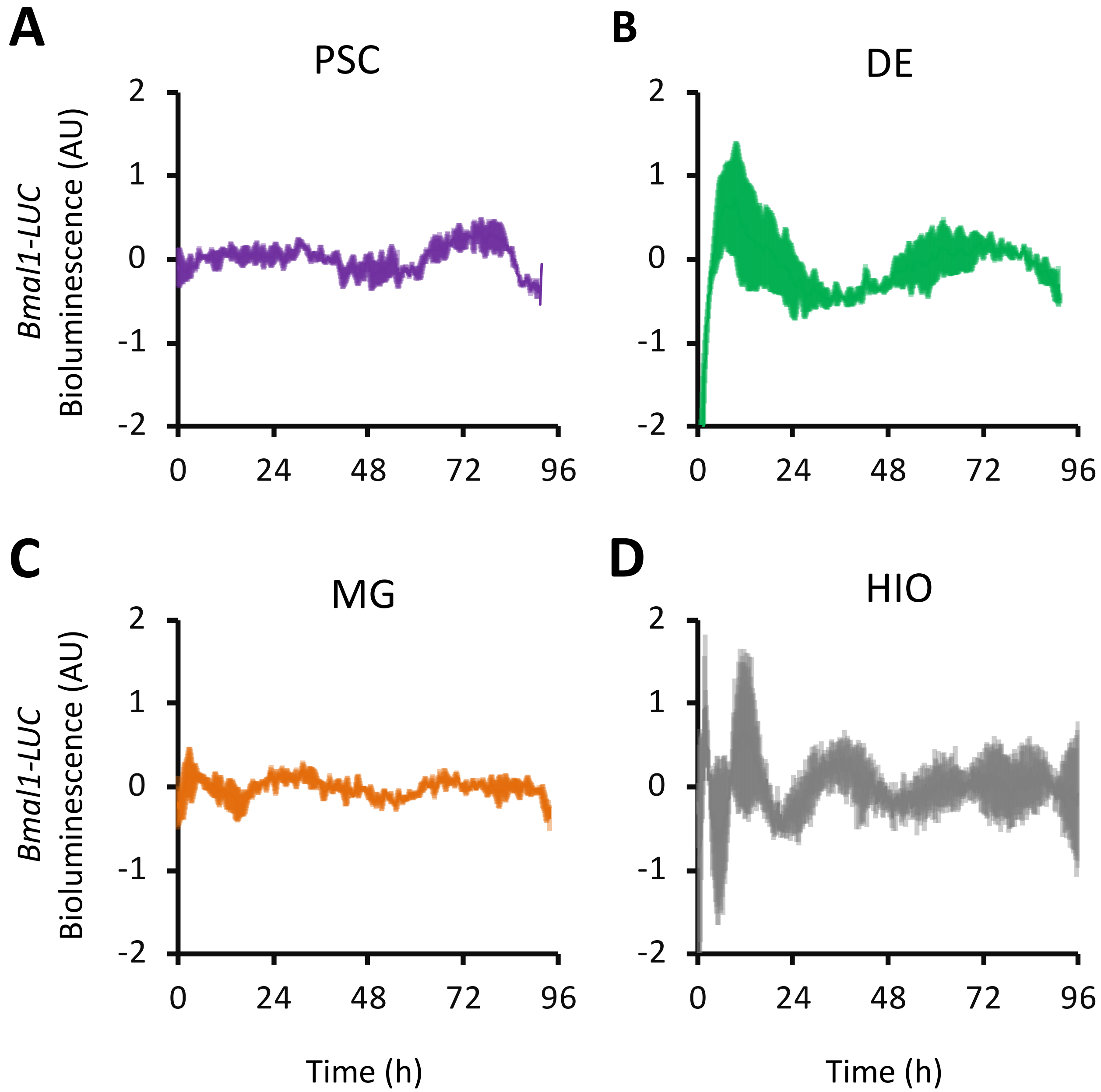
A
PSC
B
DE
C
MG
D
HIO
Bmal1-LUC
Bioluminescence (AU)
2
1
0
-1
-2
0
24
48
72
96
Time (h)

### Figure S2

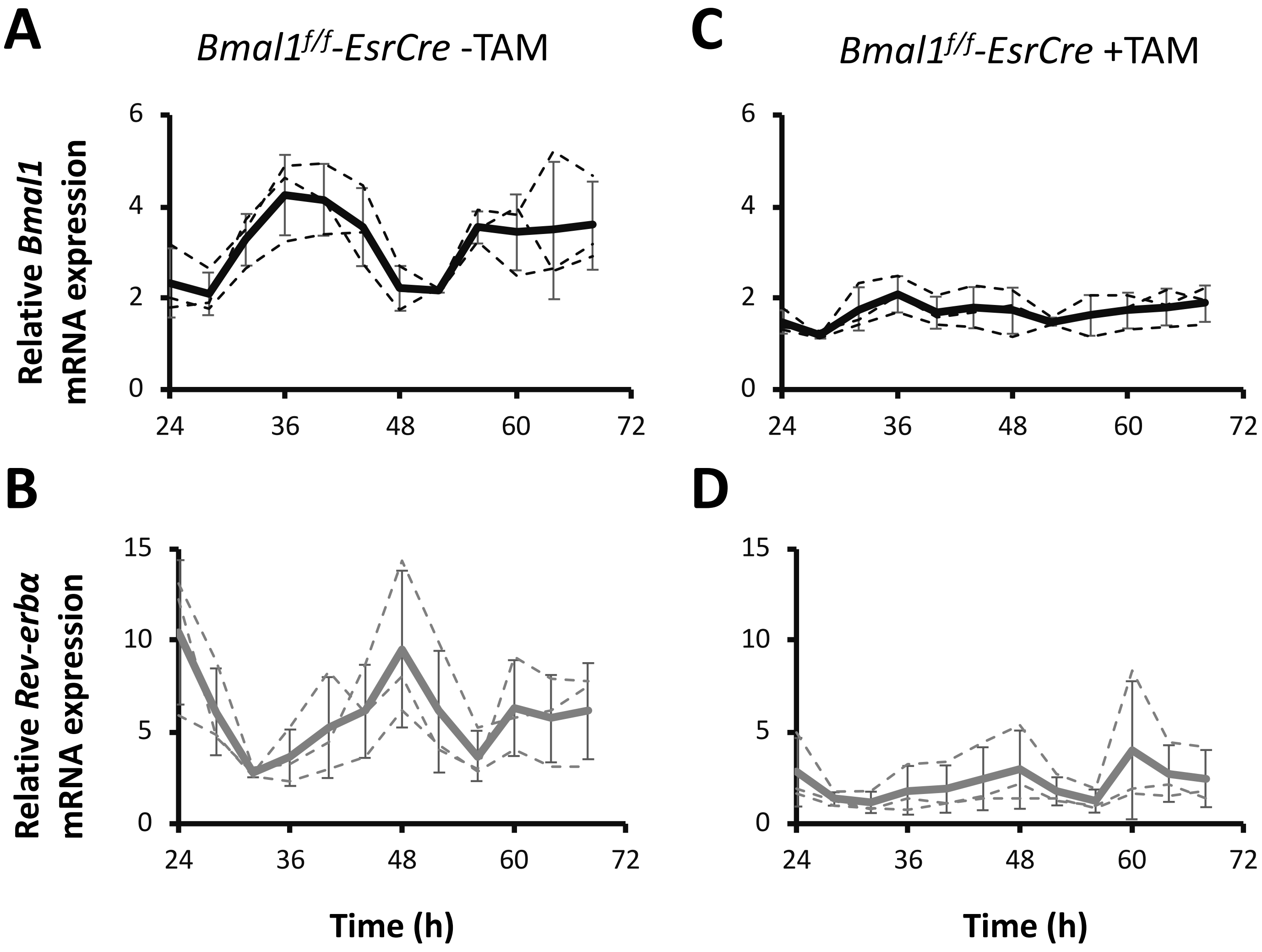
A
*Bmal1^f/f^-EsrCre* -TAM
C
*Bmal1^f/f^-EsrCre* +TAM
Relative *Bmal1* mRNA expression
6
4
2
0
24
36
48
60
72
B
D
Relative *Rev-erbα* mRNA expression
15
10
5
0
Time (h)
Time (h)

### Figure S3

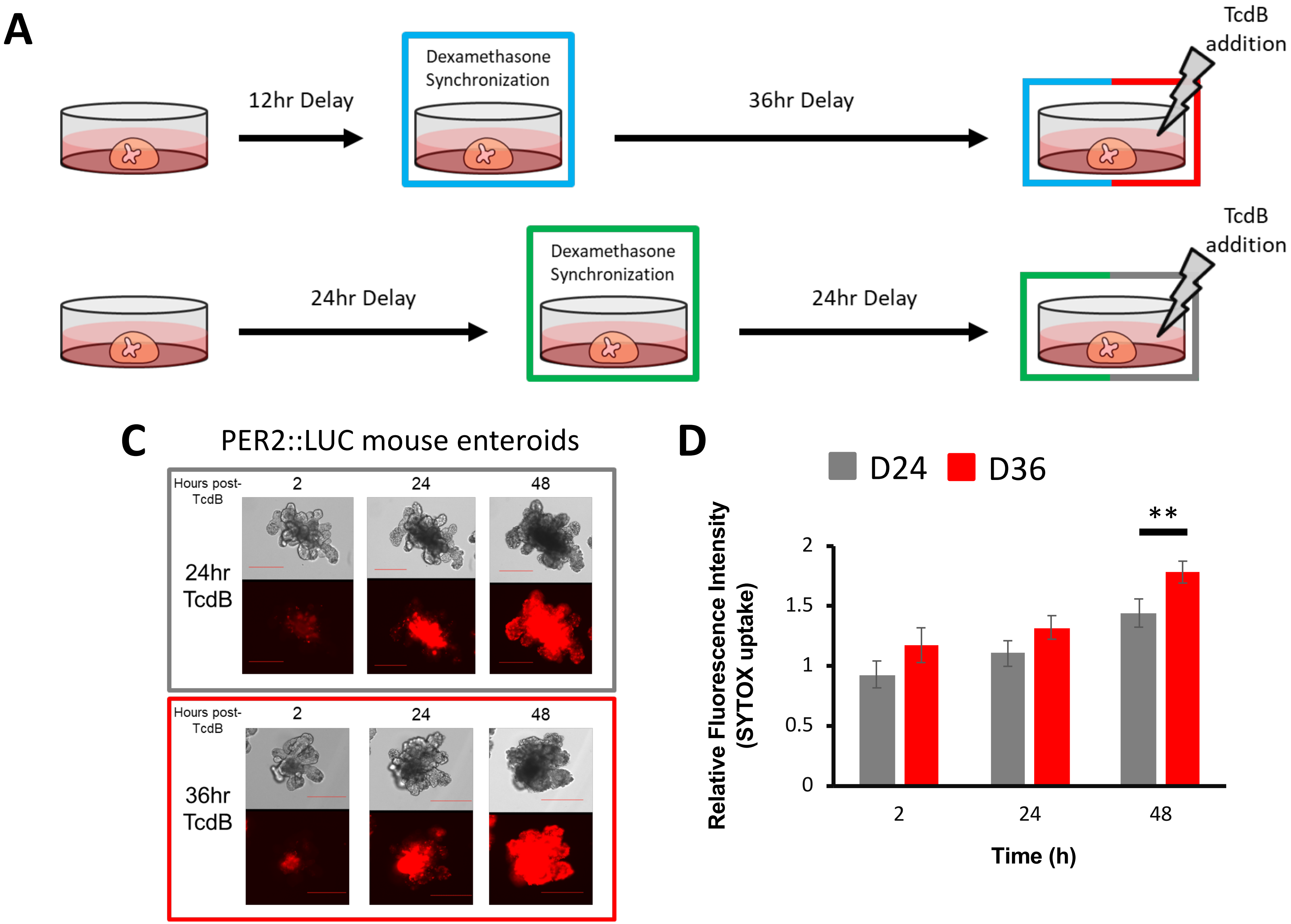

A
12hr Delay
Dexamethasone Synchronization
36hr Delay
TcdB addition
24hr Delay
Dexamethasone Synchronization
24hr Delay
TcdB addition
C
PER2::LUC mouse enteroids
Hours post-TcdB
2
24
48
24hr TcdB
Hours post-TcdB
2
24
48
36hr TcdB
D
D24
D36
**
Relative Fluorescence Intensity (SYTOX uptake)
2
1.5
1
0.5
0
2
24
48
Time (h)

### Figure S4

**A**

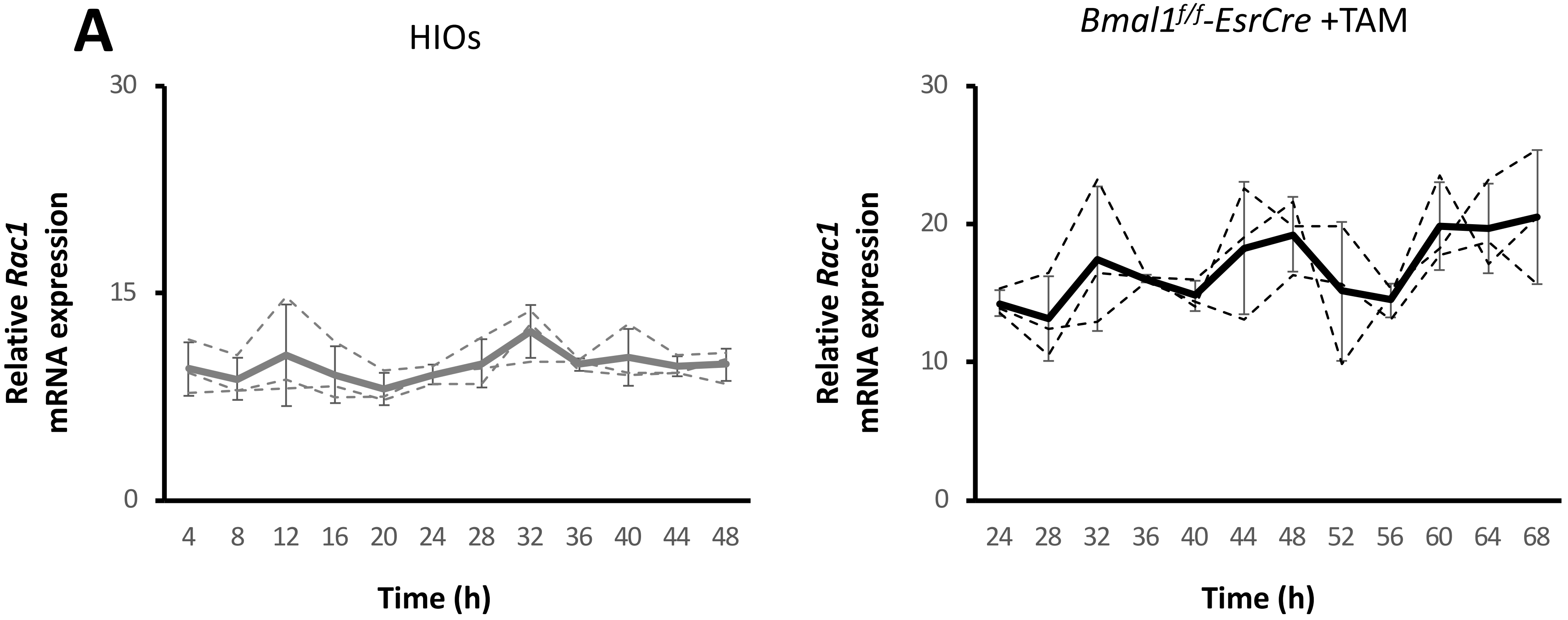
HIOs
Relative *Rac1* mRNA expression
30
15
0
4
8
12
16
20
24
28
32
36
40
44
48
Time (h)
*Bmal1$^{f/f}$-EsrCre* +TAM
Relative *Rac1* mRNA expression
30
20
10
0
24
28
32
36
40
44
48
52
56
60
64
68
Time (h)
